# Brief stimuli cast a persistent long-term trace in visual cortex

**DOI:** 10.1101/2021.02.10.430579

**Authors:** Matthias Fritsche, Samuel G. Solomon, Floris P. de Lange

## Abstract

Visual processing is strongly influenced by the recent stimulus history – a phenomenon termed adaptation. Prominent theories cast adaptation as a consequence of optimized encoding of visual information, by exploiting the temporal statistics of the world. However, this would require the visual system to track the history of individual briefly experienced events, within a stream of visual input, to build up statistical representations over longer timescales. Here, using an openly available dataset from the Allen Brain Observatory, we show that neurons in the early visual cortex of the mouse indeed maintain long-term traces of individual past stimuli that persist despite the presentation of several intervening stimuli, leading to long-term and stimulus-specific adaptation over dozens of seconds. Long-term adaptation was selectively expressed in cortical, but not in thalamic neurons, which only showed short-term adaptation. Early visual cortex thus maintains concurrent stimulus-specific memory traces of past input, enabling the visual system to build up a statistical representation of the world to optimize the encoding of new information in a changing environment.

**Significance Statement:** In the natural world, previous sensory input is predictive of current input over multi-second timescales. The visual system could exploit these predictabilities by adapting current visual processing to the long-term history of visual input. However, it is unclear whether the visual system can track the history of individual briefly experienced images, within a stream of input, to build up statistical representations over such long timescales. Here, we show that neurons in early visual cortex of the mouse brain exhibit remarkably long-term adaptation to brief stimuli, persisting over dozens of seconds, and despite the presentation of several intervening stimuli. The visual cortex thus maintains long-term traces of individual briefly experienced past images, enabling the formation of statistical representations over extended timescales.

## Introduction

Sensory processing not only depends on the current sensory input, but is influenced by the recent stimulus history. For instance, neurons in visual cortex change their responsivity and stimulus preferences following the exposure to previous visual stimuli, commonly referred to as neural adaptation (Müller et al., 1999; Dragoi et al., 2000, 2001; Kohn and Movshon, 2003, 2004). Prominent theories of adaptation posit that changes in neural responsivity can be explained by optimally efficient encoding of visual information, given temporal regularities in the recent input (Barlow, 1961; Barlow and Földiák, 1989; Weber et al., 2019). Indeed, the visual world exhibits strong temporal regularities (Dong and Atick, 1995; Simoncelli and Olshausen, 2001; Schwartz et al., 2007); for example, in natural viewing behavior, orientation information tends to be preserved across successive time points and thus stable over extended timescales (Felsen et al., 2005; van Bergen and Jehee, 2019). These temporal correlations in natural visual input can therefore be exploited by the visual system by adapting the encoding of new sensory information to the history of recent visual input. Crucially however, it is unclear over which timescales the visual system can track the history of previous input to exploit natural temporal correlations during sensory encoding.

In the natural world, previous sensory input is predictive of current input over extended timescales of multiple seconds (van Bergen and Jehee, 2019) and the visual system could exploit these predictabilities by adapting current visual processing to the long-term history of visual input. While several previous studies have indeed found evidence for long-term adaptation in early sensory cortical areas, lasting up to minutes, these studies measured neural adaptation following long stimulus presentations of dozens of seconds (Dragoi et al., 2000; Patterson et al., 2013), or in response to many brief presentations of the same stimulus (Ulanovsky, 2004; Kuravi and Vogels, 2017; Peter et al., 2020) – both reflecting very untypical sensory input under natural conditions. In contrast, neural adaptation in response to individual briefly presented stimuli has been found to be short-lived, rarely observable beyond time lags of a few hundred milliseconds in primary visual cortex of macaque monkeys and mice (Patterson et al., 2013; Jin et al., 2019; Kim et al., 2019; Jin and Glickfeld, 2020). This begs the question of whether the visual system can track the history of briefly experienced images over extended timescales, to exploit the temporal correlations present in natural input. Furthermore, it is unclear whether the visual system can maintain memory traces of the longterm history of previously experienced stimuli in the face of intervening input, or whether traces of temporally remote stimuli are eradicated by new visual inputs. Persistent memory traces, surviving the encoding of intervening visual input, would be crucial to build up robust statistical representations of the world over longer timescales.

In order to test whether neurons in early visual areas maintain long-term traces of briefly presented past stimuli, which are robust intervening visual input, we leveraged a large and unique dataset of electrophysiological recordings in the mouse visual system (Allen Brain Observatory – Neuropixels Visual Coding; Siegle et al., 2021). We characterized the recovery time course of neural adaptation in response to brief drifting and static grating stimuli across the visual system of awake mice. Neurons in the mouse primary visual cortex exhibit selectivity for orientation (Niell and Stryker, 2008; Liu et al., 2011; Tan et al., 2011), and undergo orientation-specific adaptation, tuned to the orientation difference between previous and current stimulus (Jin et al., 2019). This makes the mouse visual system suitable for probing the timescales of orientation-specific adaptation. The use of high density extracellular electrophysiology probes (Jun et al., 2017) further enabled us to study the temporal dynamics of adaptation across multiple brain areas across the visual hierarchy, in the thalamus, primary and extrastriate visual cortex. It has been previously proposed that temporal integration timescales increase along the cortical hierarchy (Hasson et al., 2008; Lerner et al., 2011; Honey et al., 2012; Murray et al., 2014). Beyond testing whether neurons in early visual areas exhibit long-term adaptation, we therefore further investigated whether a similar hierarchy of temporal dynamics may exist for stimulus-specific adaptation in the mouse visual system. Importantly, while long-term adaptation, also in the face of intervening input, has been observed in higher-order visual areas in infero-temporal cortex (McMahon and Olson, 2007), this form of adaptation appears to be task-dependent (Henson et al., 2002; Henson, 2016) and related to memory recall (Meyer and Rust, 2018). Here, we focus on the early and automatic sensory encoding of the environment, taking place in both primary and higher-order visual areas while mice viewed the stimuli passively, without an explicit task.

To preview, we found remarkably long timescales of stimulus-specific adaptation in response to brief visual stimuli in cortical visual areas, persisting over dozens of seconds, despite the presentation of several intervening stimuli. While decay of adaptation was long-lived across primary and extrastriate visual cortex, neurons in the thalamus only showed short-lived adaptation to drifting gratings, limited to the processing of temporally adjacent stimuli. Long-term adaptation in visual cortex is thus not inherited from the thalamus, and likely relies on cortical plasticity. This long-term adaptation was also evident after the exposure to more rapidly presented brief static gratings, albeit with a less clear difference in temporal decay between cortex and thalamus. This replication of long-term adaptation to briefer, more rapidly presented stimuli underlines the robustness and ecological validity of the longterm temporal dependencies. Our results indicate that early visual cortex maintains concurrent stimulus-specific memory traces of past briefly experienced input that are robust to intervening visual input. This dependence on the broader temporal context may enable the visual system to efficiently represent information in a slowly changing environment (Schwartz et al., 2007; Weber et al., 2019).

## Materials & Methods

### Dataset

All analyses were conducted on the openly available Neuropixels Visual Coding dataset of the Allen Brain Observatory (Siegle et al., 2021). This dataset surveys spiking activity from a large number of neurons across a wide variety of regions in the mouse brain, using high-density extracellular electrophysiology probes (Neuropixels silicon probes; Jun et al., 2017). Experiments were designed to study the activity of the visual cortex and thalamus in the context of passive visual stimulation. Here, we focused on a subset of experiments, termed the Brain Observatory 1.1 dataset. The Brain Observatory 1.1 dataset comprises recordings in 32 mice (16 C57BL/6J wild type mice and three transgenic lines: 6 Sst-IRES-Cre x Ai32, 5 Pvalb-IRES-Cre x Ai32 and 5 Vip-IRES-Cre x Ai32; of either sex). The three transgenic lines were included to facilitate the identification of inhibitory interneuron sub-classes using opto-tagging. For the purpose of the current research question, we analyzed the data of all 32 mice, irrespective of transgenic lines. Mice were maintained in the Allen Institute for Brain Science animal facility and used in accordance with protocols approved by the Allen Institute’s Institutional Animal Care and Use Committee. For a detailed description of the entire Neuropixels Visual Coding protocol see Siegle et al. (2021). All data are openly available through the AllenSDK (https://allensdk.readthedocs.io/en/latest/visual_coding_neuropixels.html).

### Stimuli

During Brain Observatory 1.1 experiments, mice passively viewed a variety of different stimulus types. Here, we focused on a subset of stimuli: full-field drifting and static grating stimuli (**Figures 1B** and **6A**). Visual stimuli were generated using custom scripts based on PsychoPy (Peirce, 2007) and were displayed using an ASUS PA248Q LCD monitor, with 1920 x 1200 pixels (21.93 in wide, 60 Hz refresh rate). Stimuli were presented monocularly, and the monitor was positioned 15 cm from the mouse’s right eye and spanned 120° x 95° of visual space prior to stimulus warping. Each monitor was gamma corrected and had a mean luminance of 50 cd/m^2^. To account for the close viewing angle of the mouse, a spherical warping was applied to all stimuli to ensure that the apparent size, speed, and spatial frequency were constant across the monitor as seen from the mouse’s perspective. For more details see Siegle et al. (2021).

**Figure 1.**
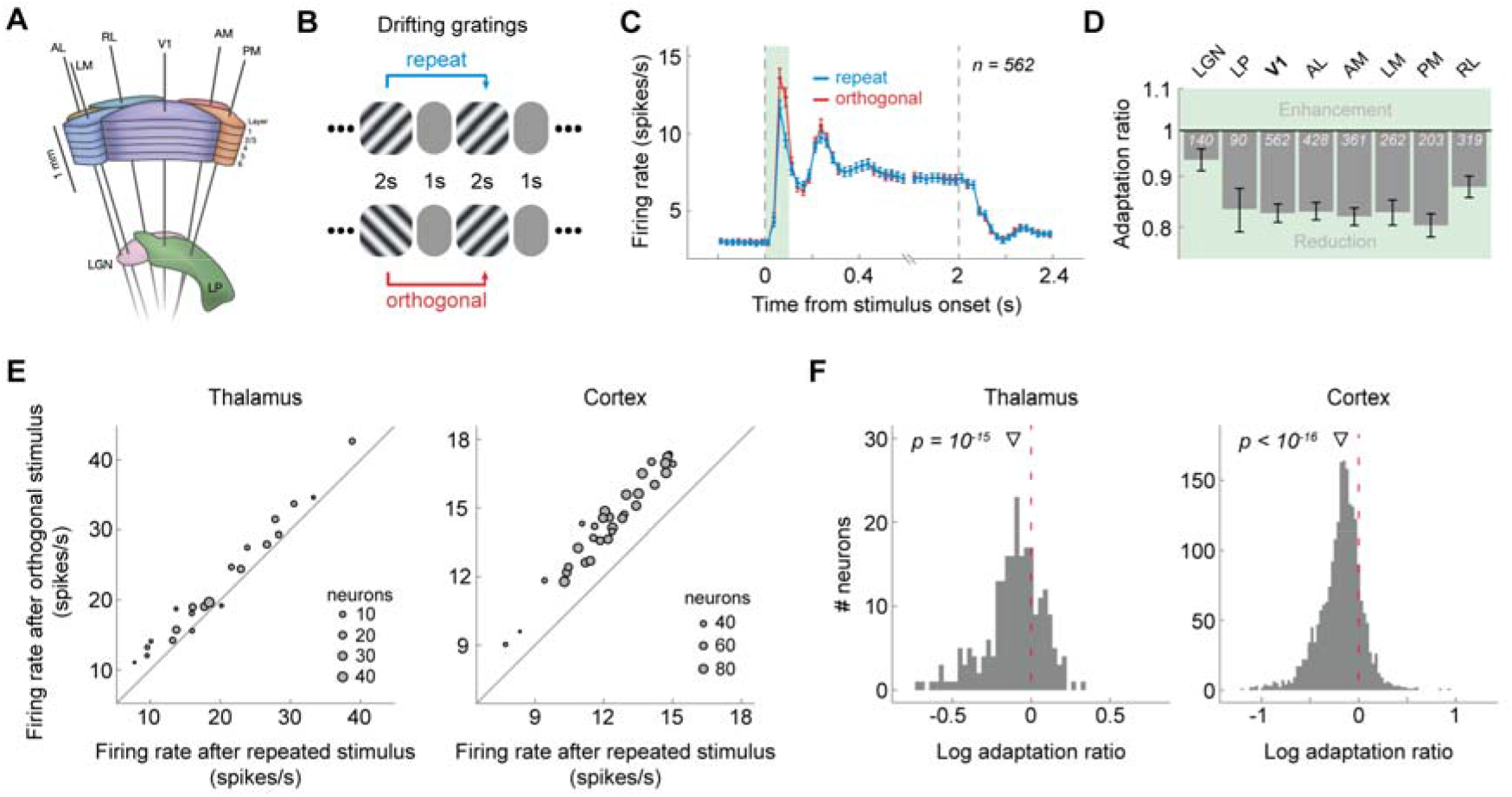
Visual cortex and thalamus exhibit orientation-specific adaptation to the immediately preceding (1-back) grating. **(A)** Schematic of Neuropixels probe insertion trajectories through visual cortical and thalamic areas (adapted from Siegle et al. (2021). **(B)** Presentation sequence of drifting grating stimuli. Mice were shown drifting gratings with a duration of 2 seconds, separated by a 1-second grey screen. Gratings were drifting in one of 8 different directions (0°, 45°, 90°, 135°, 180°, 225°, 270°, 315°) and were presented in random order. For the analysis of orientation-specific adaptation, we contrasted activity to gratings preceded by gratings of the same orientation (*repeat*, blue) with that elicited by gratings preceded by a grating of the orthogonal orientation (*orthogonal*, red). **(C)** Population peristimulus time histograms of neurons in V1 for *repeat* and *orthogonal* conditions. The transient response is reduced when the same orientation is successively repeated, indicating orientation-specific adaptation. Subsequent analyses focused on this transient response (0 – 100 ms, green shaded area). Vertical dashed lines denote stimulus onset and offset, respectively. Binwidth = 25 ms. Error bars show *SEMs*. **(D)** 1-back adaptation ratios of transient responses across visual areas. Adaptation ratios were computed by dividing each neuron’s firing rate for *repeat* by that for *orthogonal* stimulus presentations and therefore express the response magnitude to a repeated stimulus orientation relative to that elicited by the same stimulus orientation, but preceded by a grating with the orthogonal orientation. Adaptation ratios smaller than 1 indicate adaptation. All visual areas show significant 1-back adaptation. Error bars denote bootstrapped 95% confidence intervals. White numbers indicate the number of neurons in each area. **(E)** The average firing rate to a stimulus preceded by a stimulus with the same orientation (*x*-axis) is consistently smaller than the firing rate to a stimulus preceded by a stimulus with the orthogonal orientation (*y*-axis) across mice (grey dots denote different mice; size scaled by the number of neurons of each mouse) in both thalamus (left) and cortex (right), as indicated by datapoints positioned above the diagonal. **(F)** Histograms of singleneuron adaptation ratios (log-transformed) in thalamus (left) and cortex (right). Negative *x*-values indicate adaptation and the red dashed line marks zero adaptation (i.e., equal firing rates for repeat and orthogonal conditions). The triangle shape indicates the mean adaptation across the population of neurons with p-value indicating the significance of the population mean. List of acronyms: Dorsolateral geniculate nucleus of the thalamus (*LGN*), latero-posterior nucleus of the thalamus (*LP*), primary visual cortex (*V1*), antero-lateral area (*AL*), antero-medial area (*AM*), latero-medial area (*LM*), postero-medial area (*PM*), rostro-lateral area (*RL*).

**Figure 2.**
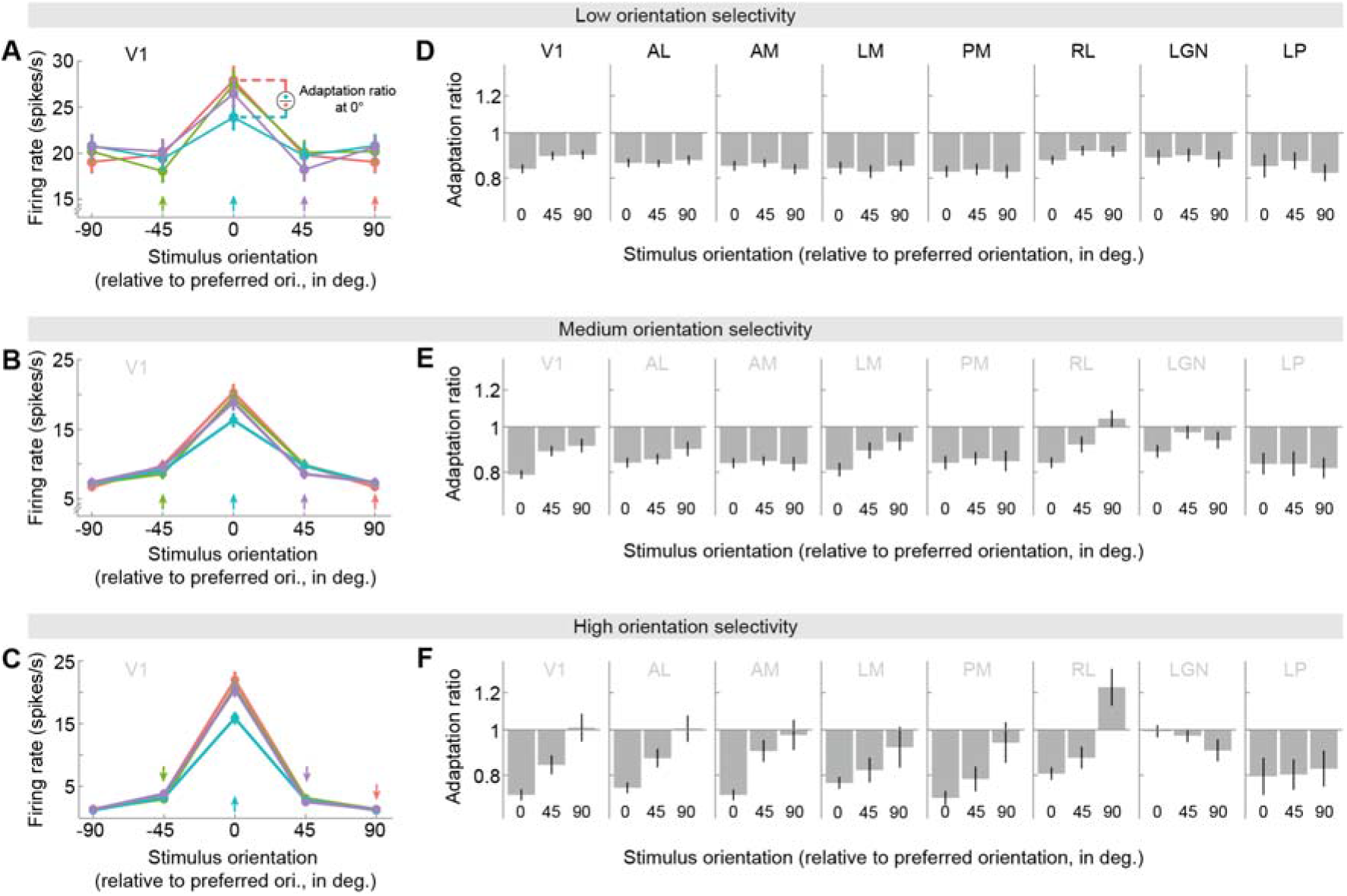
Adaptation depends on orientation tuning and adaptor/test orientation. **(A, B, C)** Orientation tuning curves in V1 for units of low **(A)**, medium **(B)** or high **(C)** orientation selectivity (tertile split, see **Materials & Methods**), following adaptation to different 1-back grating orientations (colored arrows). Stimulus and adaptor orientations are expressed relative to each neuron’s preferred orientation. Tuning curves show local response reductions to the adapted orientation. **(D, E, F)** Adaptation ratios as a function of the adaptor and test orientation relative to the neuron’s preferred orientation. For instance, the adaptation ratio for a relative stimulus orientation of 0° compares the visual response to a test grating with the neuron’s preferred orientation, when it is preceded by an adaptor grating with the same (preferred) orientation, versus when it is preceded by the orthogonal (non-preferred) adaptor orientation (see illustration in panel **A**). In V1 (panels **D**, **E** and **F**, leftmost columns), adaptation was strongest when adaptor and test stimuli corresponded to the preferred orientation of the neuron, and decreased when adapting and testing with less preferred orientations (significant main effect of relative orientation, p = 4e-11). This relationship was particularly strong in neurons exhibiting high orientation selectivity (significant interaction between relative adaptor/test orientation and orientation selectivity, p = 0.005; for definition of orientation selectivity see **Materials & Methods**). Nevertheless, there was clear adaptation for all adaptor orientations as indicated by 1-back adaptation ratios consistently smaller than 1 (all p < 0.004, corrected for multiple comparisons), except for non-preferred (90°) adaptor and test stimuli of highly selective units (panel **F**, leftmost column, 90°, p = 0.88). This overall pattern of adaptation effects was qualitatively similar across cortical visual areas (panels **D**, **E** and **F**, columns 2 to 5). In thalamic areas (panels **D**, **E** and **F**, two rightmost columns), there was no evidence for a dependence of adaptation on orientation preference (no significant main effects of relative adaptor/test orientation: LGN, p = 0.28; LP, p = 0.91; no significant interactions between relative adaptor/test orientation and orientation selectivity: LGN, p = 0.24; LP, p = 0.92), likely due to the overall lower degree of orientation selectivity of thalamic neurons.

**Figure 3.**
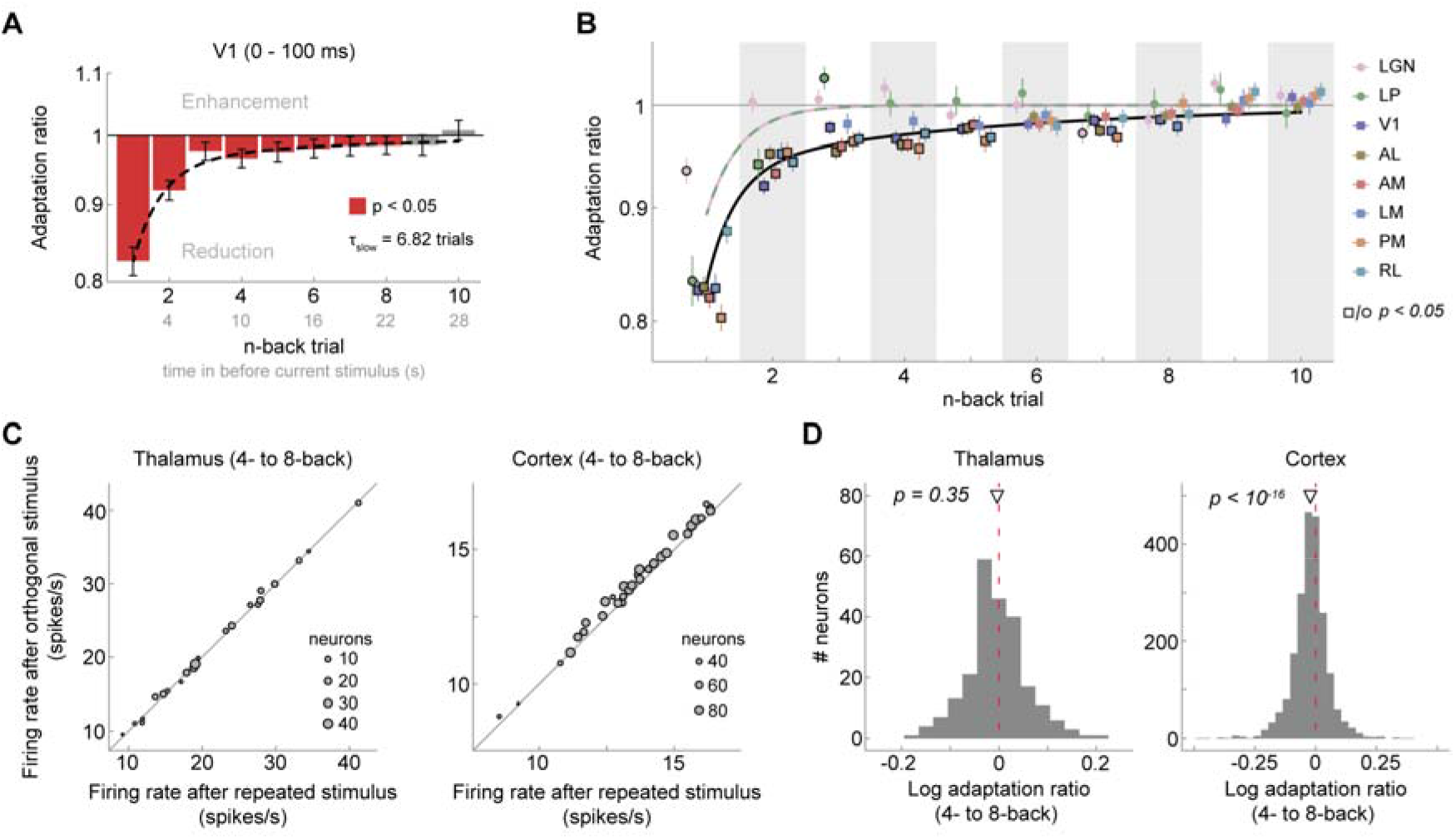
Visual cortex, but not thalamus, exhibits long-term adaptation. **(A)** Adaptation ratios of neurons in V1 as a function of the n-back trial. Strongest adaptation occurred in response to the 1-back stimulus, but stimuli encountered up to 8 presentations in the past (seen 22 seconds ago) still exerted significant adaptation effect on the current visual response, despite the presentation of intervening stimuli (red bars, p < 0.05, corrected for multiple comparisons). The decay of adaptation over n-back trials was well captured by a double-exponential decay model with a fast- and slow-decaying adaptation component (black dashed line; *a_fast_* = 13.99%, t_fast_ = 0.85 trials, *a_slow_* = 3.45%, t_slow_ = 6.82 trials). Error bars denote bootstrapped 95% confidence intervals. **(B)** Adaptation ratio as function of n-back trial for different visual areas (color-coded). While adaptation decays similarly and slowly across cortical visual areas (square symbols), and is generally significant for up to 6-8 trials back (symbols with black border, p < 0.05, corrected for multiple comparisons per area), it decays more rapidly in thalamic areas LGN and LP (circle symbols). Black and lilac-green lines illustrate the best fitting exponential decay models for cortex and thalamus. Error bars denote standard errors of the mean. **(C)** Average firing rates per mouse when the 4- to 8-back orientation was repeated (*x*-axis) or orthogonal (*y*-axis) relative to the current orientation. Mice exhibit consistent long-term adaptation in cortex (right) but not in thalamus (left). **(D)** Histograms of single-neuron adaptation ratios (log-transformed) in thalamus (left) and cortex (right).

**Figure 4.**
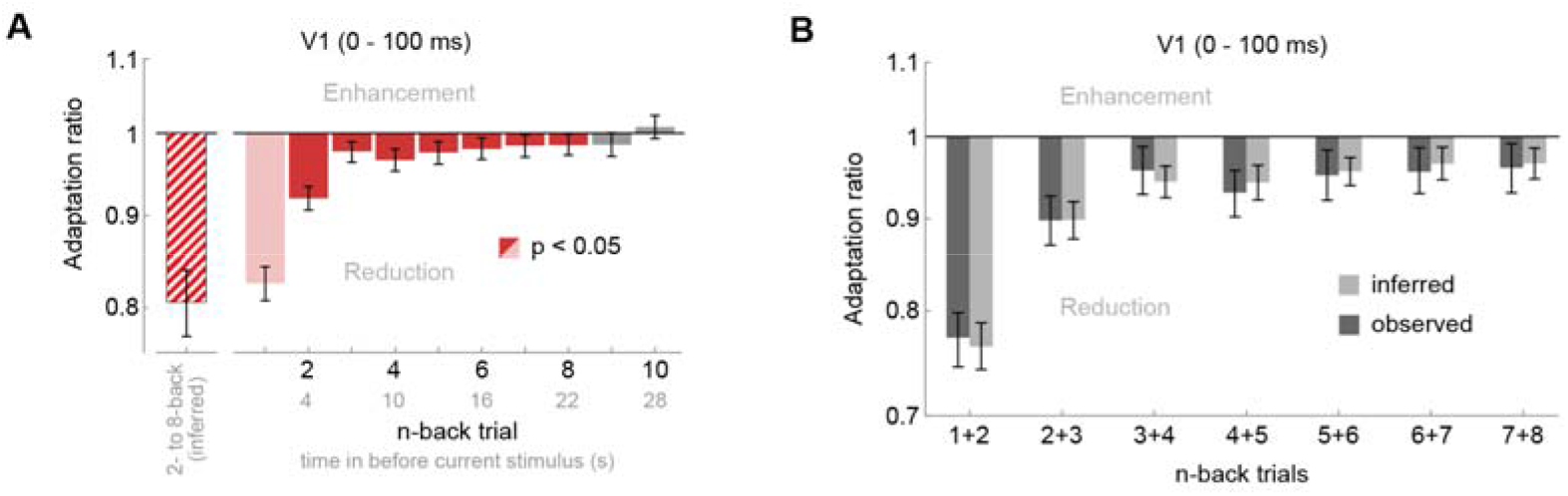
Cumulative adaptation effects in V1. Random sequences of grating orientations, as the ones used in the current experiment, prevent any systematic accumulation of adaptation across multiple stimulus presentations. While this allows us to study the influence of individual n-back stimuli on the current visual response, it underestimates the influence of long-term adaptation in natural environments, in which orientations tend to remain stable over prolonged time periods (van Bergen & Jehee, 2019), therefore leading to an accumulation of adaptation. Panel **(A)** serves to illustrate that the adaptation effects of 2- to 8-back stimuli (red bars), albeit small when taken individually, together may lead to a considerable reduction of the current response (19% reduction; red-striped bar) that even outweighs the adaptation effect of the 1-back stimulus (17% reduction; light red bar). Importantly, the cumulative influence of repeating 2- to 8-back grating orientations could not be estimated empirically in the current dataset, since such streaks of orientation repetitions are exceedingly rare for random sequences (probability of ~0.006%). Here, we inferred the cumulative response reduction by assuming that the adaptation effects of previous stimuli accumulate approximately linearly. The inferred cumulative adaptation ratio was then calculated as 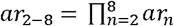. where *ar_2-8_* is the cumulative adaptation ratio of 2- to 8-back stimuli, and *ar_n_* denotes the empirically estimated adaptation ratio of an individual n-back stimulus. **(B)** To evaluate whether the assumption of a linear accumulation of adaptation approximately holds, we compared the empirically observed adaptation effect when two previous adjacent stimuli had the same orientation as the current stimulus (dark grey bars; ~6.25% of all trials), to the cumulative adaptation effect inferred from individual n-back adaptation estimates (light grey bars). The empirically observed adaptation effect of two successive stimuli roughly matched the predicted adaptation effect, suggesting that adaptation accumulates approximately linearly in the current setting. All error bars denote 95% CIs.

**Figure 5.**
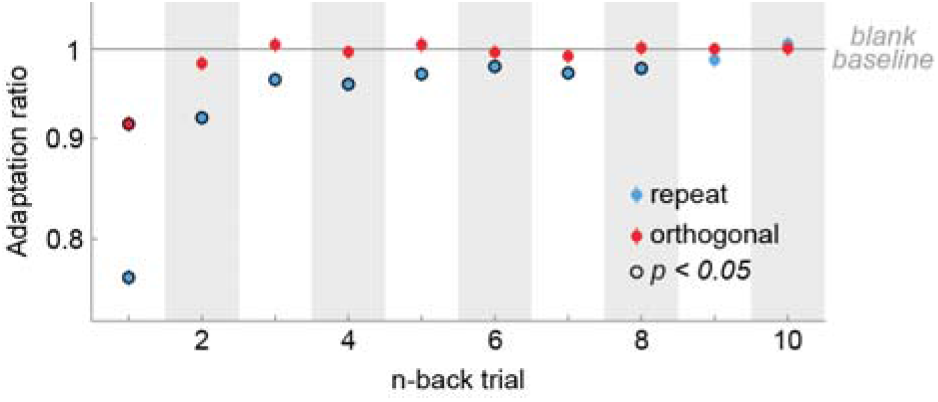
Cortical long-term adaptation is driven by repeated stimulus orientations. We expressed the response modulation of neurons across all cortical areas by n-back repeated and orthogonal trials relative to a neutral baseline, in which no stimulus was presented on the n-back trial. To this end, we computed adaptation ratios by dividing each neuron’s firing rate for *repeat* stimulus presentations by that of *blank* stimulus presentations (blue data points), or *orthogonal* divided by *blank* stimulus presentations (red data points). While the suppressive effects of orthogonal stimuli decays quickly, repeated stimuli exert long-term suppression for up to 8 trials. Error bars denote bootstrapped 95% confidence intervals.

**Figure 6.**
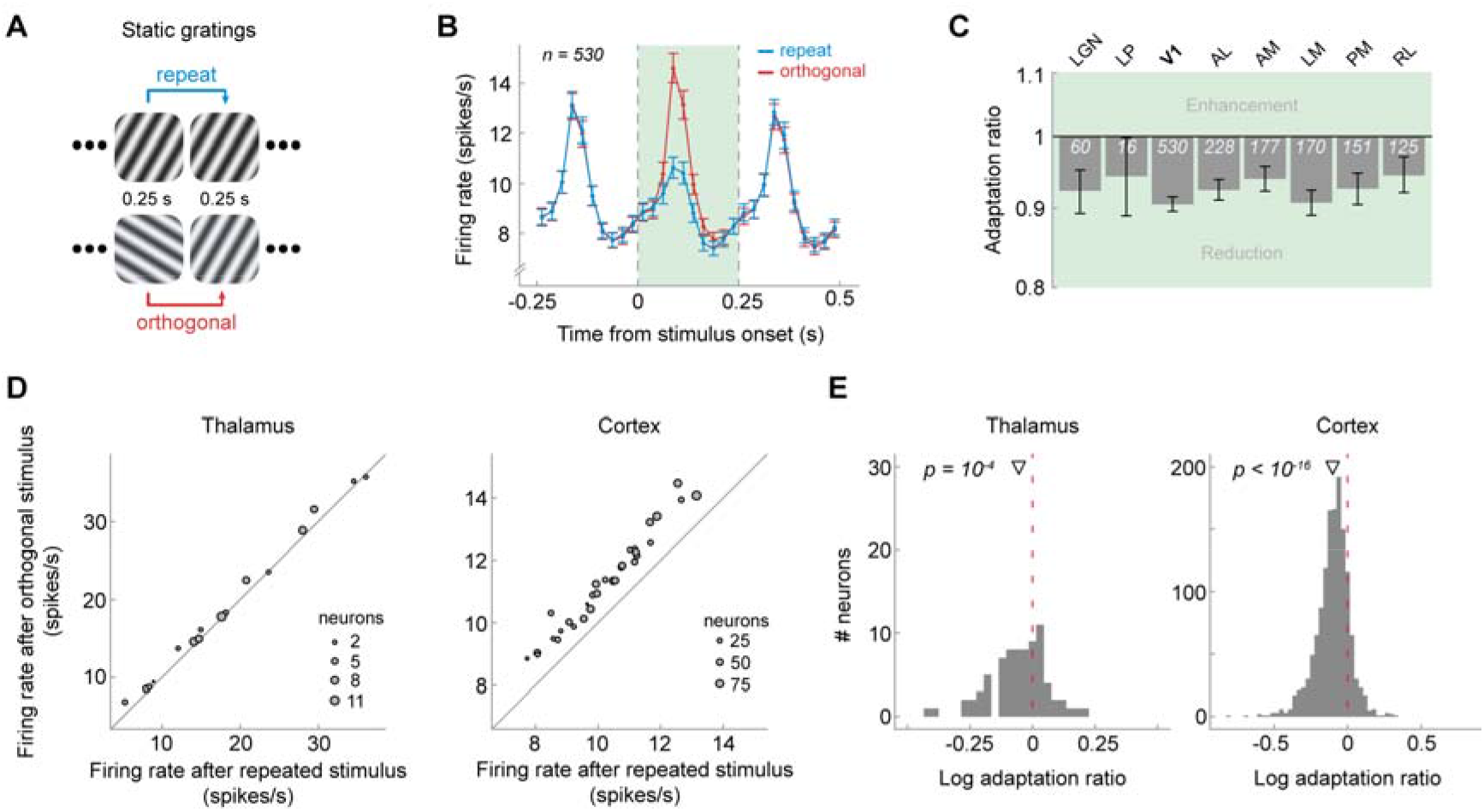
Visual cortex exhibits adaptation in response t*o* immediately preceding briefly presented static gratings. **(A)** Presentation sequence of static grating stimuli. Mice were shown static gratings with a duration of 250 ms with no intervening grey period. Gratings had one of six orientations (0°, 30°, 60°, 90°, 120°, 150°), five spatial frequencies (0.02, 0.04, 0.08, 0.16, 0.32 cycles/°), and four phases (0, 0.25, 0.5, 0.75). The order of grating presentations was randomized. Similar to the analysis of drifting gratings, we contrasted activity to gratings preceded by gratings of the same orientation (*repeat*, blue) with that elicited by gratings preceded by a grating of the orthogonal orientation (*orthogonal*, red). **(B)** Population peristimulus time histograms of neurons in V1 for *repeat* and *orthogonal* conditions. The visual response to the current stimulus (green shaded area) was reduced when the previous stimulus had the same orientation as the current stimulus (*repeat*), indicating orientation-specific adaptation. Vertical dashed lines denote onset and offset of the current stimulus, respectively. Binwidth = 25 ms. Error bars show *SEMs*. **(C)** 1-back adaptation ratios across visual areas. All areas show significant 1-back adaptation. Error bars denote bootstrapped 95% confidence intervals. White numbers indicate the number of neurons in each area. **(D)** Mice show consistently reduced firing rates after a repeated versus orthogonal orientation, as indicated by datapoints falling above the diagonal. Same conventions as in **Fig. 1E**. **(E)** Histograms of singleneuron adaptation ratios (log-transformed) in thalamus (left) and cortex (right).

Full-field drifting gratings were shown with a spatial frequency of 0.04 cycles/deg, 80% contrast, 8 directions (0°, 45°, 90°, 135°, 180°, 225°, 270°, 315°, clockwise from 0° = right-to-left) and 5 temporal frequencies (1, 2, 4, 8, and 15 Hz), with 15 repeats per condition, resulting in a total number of 600 drifting grating presentations, divided across three blocks. Drifting gratings were presented for 2 seconds, followed by a 1 second inter-stimulus interval (grey screen). Gratings of different directions and temporal frequencies were presented in random order and were interleaved by the presentation of 30 blank trials, in which only a grey screen was shown.

Static gratings were shown at 6 different orientations (0°, 30°, 60°, 90°, 120°, 150°, clockwise from 0° = vertical), 5 spatial frequencies (0.02, 0.04, 0.08, 0.16, 0.32 cycles/degree), and 4 phases (0, 0.25, 0.5, 0.75). They were presented for 0.25 seconds, with no intervening grey period. Gratings with each combination of orientation, spatial frequency, and phase were presented ~50 times in a random order, resulting in a total of 6000 grating presentations, divided across three blocks. There were blank sweeps (i.e. mean luminance grey instead of grating) presented roughly once every 25 gratings.

### Data analyses

All data analyses were performed using custom code written in Python, Matlab and R. All code will be made openly available on the Donders Institute for Brain, Cognition and Behavior repository at https://data.donders.ru.nl.

#### Unit exclusion

To filter out units (i.e. putative neurons) that were likely to be highly contaminated or missing lots of spikes, we applied the default quality metrics of the AllenSDK. This entailed excluding units with ISI violations larger than 0.5 (Hill et al., 2011), an amplitude cutoff larger than 0.1 and a presence ratio smaller than 0.9 (for more details see https://allensdk.readthedocs.io/en/latest/_static/examples/nb/ecephys_quality_metrics.html). For the analysis of drifting gratings, we defined visually responsive units as those units whose average firing rate during the first 100 ms of stimulus presentation of the unit’s preferred orientation (eliciting the highest firing rate) was larger than 5 Hz and larger than 1 *SD* of the firing rate during the first 100 ms of grey screen presentations. For the analyses of static gratings, we applied the same inclusion criteria, but computed firing rates over the whole stimulus duration (i.e. 250 ms). We chose to use a longer time window for analyzing static grating adaptation, since due to the back-to-back presentation of the static gratings, visual responses to the previous stimulus overlapped with the initial time window of the current stimulus, thereby increasing response variability in this early time window. However, largely similar results were obtained when performing the analyses on the same time window used in the drifting grating experiment (0 to 100 ms). In order to assess whether the choice of the minimum firing rate threshold of 5 Hz had a substantial impact on our results, we repeated the analyses with a more conservative (10 Hz) and less conservative (2.5 Hz) threshold, but obtained qualitatively similar results. In our further investigation of sensory adaptation, we focused on those regions that contained a minimum of 50 visually responsive units (for an overview of included regions and unit counts per region see **Figures 1D and 6C**). All subsequent analyses were performed on visually responsive units only.

#### Orientation-specific adaptation to drifting gratings

To investigate orientation-specific adaptation, for each unit we compared firing rates in response to a current grating when this grating was preceded by a grating with the same orientation (*repeat*) or by its orthogonal orientation (*orthogonal*), irrespective of the temporal frequencies of current and previous gratings. Note that repeat trial pairs could consist of gratings with opposite drifting directions, but with the same orientation. Investigating orientation-rather than direction-specific adaptation had the advantage of maximizing the number of repeat and orthogonal trial pairs occurring across the random trialsequence. Adaptation was quantified in the form of an adaptation ratio:

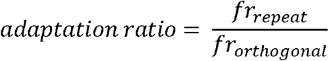

where *fr_repeat_* and *fr_orthogonal_*, are the firing rates in response to a repeated and orthogonal stimulus, respectively. The adaptation ratio expresses the response magnitude to a repeated stimulus orientation relative to that elicited by the same stimulus orientation, but preceded by a grating with the orthogonal orientation. Adaptation ratios smaller than 1 indicate a relative response reduction for orientation repetitions. Importantly, this analysis quantifies orientation-specific adaptation, as the stimulus features between repeat and orthogonal condition are the same (on average), with the only difference being the relative orientation of the adaptor stimulus. An initial exploratory analysis in one mouse suggested strongest adaptation effects for the early transient response (0-100 ms from stimulus onset). Therefore, we limited our analysis to this time window. This analysis choice was made while blind to adaptation effects beyond the 1-back grating, which were of main interest to the current study. Adaptation induced by n-back gratings was quantified in a similar manner as described above, by conditioning the data on the orientation difference (*repeat* or *orthogonal*) between the current and n-back gratings. For each region, we statistically compared log-transformed adaptation ratios of 1- to 10-back gratings to zero (indicating no adaptation) using two-tailed t-tests, while controlling the false discovery rate at an alpha-level of 0.05 using the Benjamini-Hochberg procedure.

In order to quantify the recovery time course of adaptation, we fitted exponential decay models to the 1- to 50-back adaptation ratios of each region. The recovery of adaptation in cortical areas was significantly better fit by a double exponential, compared to a single-exponential decay model, with a fast and slow decay component, of the form:

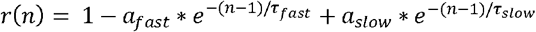

where r(n) denotes the adaptation ratio conditioned on the n-back stimulus orientation, *a_fast_, τ_fast_, a_slow_* and *τ_slow_* determine the magnitude and recovery time of the fast and slow adaptation component, respectively. Adaptation in thalamic regions was more parsimoniously explained by a single-exponential decay model of the form:

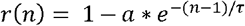

For each region, we statistically compared single- and double exponential decay models with an F-test. We used an F-test as the two decay models are nested – the single-exponential decay model is a restricted version of the double-exponential decay model. Since adaptation ratios were not normally distributed, all models were fit to log-transformed adaptation ratios by analogously log-transforming model predictions. We obtained the 95% confidence intervals of the parameter estimates with a bootstrapping procedure. In particular, for each region we resampled units with replacement and refitted the exponential decay model. We repeated this procedure 1,000 times and recorded the resulting parameter estimates of the bootstrapped sample. The 95% confidence interval was taken as the 2.5 and 97.5 percentile of the bootstrapped parameter distribution. We restricted parameter values to a wide range of plausible values (*a_fast_* = *a_slow_* = [-Inf, 0.5], *τ_fast_* = *τ_slow_* = [-50, 50]), and discarded bootstrapped estimates which lay on the boundary of the parameter range, indicating implausible fits (0.3% of bootstrapped fits).

Additionally, we investigated to which degree 1-back adaptation was dependent on the relationship between a unit’s orientation preference and the adaptor/test orientation. For instance, one may expect strongest adaptation when the repeated stimuli match the unit’s preferred orientation, due to the strong response during the adaptation period. To shed light on this question, we first binned units into three equally sized subgroups per region, based on their orientation selectivity. Orientation selectivity was quantified as

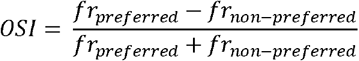

where *fr_preferred_* and *fr_non-preferred_* refer to the unit’s firing rates to its preferred orientation (eliciting the highest average firing rate) and the orthogonal orientation, respectively. The OSI ranges from 0 to 1, where 0 indicates no selectivity (identical firing rates to preferred and non-preferred orientations) and 1 indicates maximal selectivity (zero firing rate to non-preferred orientation). Subsequently, for each subgroup of units we computed adaptation ratios as a function of the previous (adaptor) and current (test) stimulus orientation relative to the unit’s preferred orientation (see **Figure 2**). To statistically test the influence of the relative adaptor/test stimulus orientation on adaptation ratios, and to test whether this influence depended on the degree of orientation selectivity of the units, we conducted a 3 x 3 mixed ANOVA, with repeated measures factor “relative adaptor/test orientation” (0, 45 and 90°) and between-unit factor “orientation selectivity” (low, medium and high OSI).

Since we found that adaptation was indeed strongest when the repeated orientations matched the unit’s preferred orientation, we repeated our analysis of the recovery time course of adaptation for these trial types. That is, we computed adaptation ratios on a subset of trials, for which the current orientation matched the unit’s preferred orientation and the previous orientation either matched (*repeat*) or was orthogonal (*orthogonal*) to the preferred orientation. While this approach had the advantage of quantifying adaptation to the most effective adaptor stimulus, it had the disadvantage of limiting the analysis to a much smaller set of trials compared to computing adaptation for all orientations. We did not observe qualitative differences between the two analysis approaches.

#### Dissociating adaptation to repeated and orthogonal drifting gratings

Thus far, we have quantified adaptation as the ratio between responses to repeated and orthogonal stimulus orientations. This analysis does not reveal whether adaptation effects are due to suppression of response when the current orientation matches that of past orientations, facilitation of response when the current orientation is orthogonal to past orientations, or a mixture of the two. The stimulus set included randomly interspersed trials during which no stimulus was presented, so we repeated the analysis described above, but quantified adaptation by comparing responses when a stimulus preceded the current trial, with responses when no stimulus was presented in the preceding trial. We computed two sets of adaptation ratios: (1) the ratio between visual responses when the n-back stimulus had the same orientation as the current stimulus (*n-back repeat*) and trials in which no stimulus was presented at the same n-back position (*n-back blank trial*). (2) the ratio between visual responses when the n-back stimulus was orthogonal to the current stimulus (*n-back orthogonal*) and *n-back blank* trials. Since blank trials were much less frequent than repeat and orthogonal trials (30 blank trials vs ~150 repeat/orthogonal trials), for these analyses we randomly subsampled repeat and orthogonal trials to match them to the lower number of blank trials.

#### Orientation-specific adaptation to static gratings

Analyses of adaptation to static gratings were similar to the analysis of the drifting grating data, with two exceptions. First, we quantified adaptation based on neural responses during the entire stimulus presentation period (0-250 ms). As discussed above, we chose to use a longer time window for analyzing static grating adaptation, since due to the back-to-back presentation of the static gratings, visual responses to the previous stimulus overlapped with the initial time window of the current stimulus, thereby increasing response variability in this early time window. However, largely similar results were obtained when performing the analyses on the same time window used in the drifting grating experiment (0 to 100 ms). Second, we only analyzed adaptation to all orientations, regardless of the units’ orientation preferences. This analysis was similar to the main analysis of drifting grating adaptation described above. Due to the rapid presentation of the static gratings, without intervening grey periods, responses persisted into the presentation period of the next grating. Since sub-selecting data according the units’ preferred orientation led to response differences in the adaptation period (i.e. larger response to preferred than orthogonal adaptor), the bleeding of the previous response into the current stimulus time window strongly biased the response to the current grating, thereby confounding genuine adaptation-induced changes in the response to the current grating. Conversely, when analyzing adaptation for all orientations, regardless of the units’ orientation tuning, the relationship between the adaptor orientations and the units’ preferred orientations were balanced across *repeat* and *orthogonal* trials, and therefore did not bias the analysis of the current response.

## Results

### Orientation-specific adaptation in visual cortex and thalamus

To investigate orientation-specific adaptation in the mouse visual system, we analyzed responses from a total of 2,365 visually responsive neurons in the visual cortex and thalamus of 32 mice (**Figure 1A**), while they were presented with sequences of drifting gratings (**Figure 1B**). We separately analyzed visual responses to gratings that were preceded by a grating of the same orientation (*repeat*) or orthogonal orientation (*orthogonal*). We found that the immediate repetition of stimulus orientation led to a marked, orientation-specific reduction in spiking activity in primary visual cortex (V1), predominantly during the early visual response (0 – 100 ms from stimulus onset, *n* = 562; **Figure 1C**, green shaded area). We quantified this orientation-specific adaptation of the transient visual response by calculating the response to a repeated orientation, relative to that following the orthogonal orientation (*1-back adaptation ratio*, see **Materials & Methods**). Adaptation reduced the response by 17% in V1 (1-back adaptation ratio: 0.83, p = 4e-57, 95% CI [0.81, 0.84]), and had similar impact in higher-level extrastriate visual areas (**Figure 1D**; 1-back adaptation ratios between 0.80 and 0.88, all p < 2e-21, two-sided t-tests, corrected for multiple comparisons). We also found orientation-specific adaptation in the dorsolateral geniculate nucleus of the thalamus (LGN, *n* = 140; 1-back adaptation ratio: 0.93, p = 8e-7, 95% CI [0.91, 0.96]), and the lateral posterior nucleus of the thalamus (LP, *n* = 90; 1-back adaptation ratio: 0.83, p = 3e-10, 95% CI [0.79, 0.88]). Of note, in this analysis, we focus on *stimulus-specific* adaptation, sensitive to the orientation difference between previous and current stimulus (repeat versus orthogonal). This analysis is not sensitive to additional untuned adaptation effects, which occur in response to previous stimuli of any orientation and thus do not track the history of previous orientations (see **Figure 5** for a complementary analysis quantifying adaptation to repeat and orthogonal stimuli, separately, versus adaptation in response to a blank grey screen). This may explain why the current response reductions are slightly smaller than previous reports of adaptation that comprise both orientation-specific and unspecific adaptation (Patterson et al., 2013; Jin et al., 2019; Jin and Glickfeld, 2020). The orientation-specific response reductions for immediate stimulus repetition were highly consistent across mice (**Figure 1E**). While 1-back adaptation was generally strongest when neurons were tested with their preferred orientation, neurons also showed robust orientation selective adaptation when probed at non-preferred orientations (**Figure 2**). In our subsequent analyses of long-term adaptation, we therefore averaged adaptation across all stimulus orientations, regardless of the neurons’ orientation preference, but qualitatively similar results were obtained when only considering trials in which stimuli matched a neurons’ preferred orientation. Overall, these findings indicate robust orientation-specific adaptation of neurons in visual cortex and thalamus to gratings presented in the immediate past.

### Long-term adaptation in visual cortex but not in thalamus

In order to investigate the timescale over which adaptation influences subsequent visual processing, we computed adaptation ratios based on the orientation difference (i.e., repeat versus orthogonal) between the current grating, and gratings at different n-back timepoints. Surprisingly, we found that neurons in V1 exhibited significant adaptation effects to stimuli seen up to 8 presentations (or 22 seconds) in the past, despite the presentation of multiple intervening stimuli (**Figure 3A**). It is worth noting that although individual past stimuli had subtle effects on the current response, cumulative adaptation to the remote stimulus history outweighed the immediate adaptation effect (17% response reduction to 1-back stimulus vs. 19% cumulative response reduction to 2- to 8-back stimuli; **Figure 4**). In natural temporally correlated environments, long-term adaptation may thus have even greater weight than immediate adaptation effects. Therefore, the joint long-term stimulus history exerts a considerable influence on current sensory processing.

In contrast to adaptation in cortex, adaptation in the thalamus appeared to be limited to the 1-back (LGN) or 2-back trial (LP; **Figure 3B**). Indeed, adaptation to temporally remote stimuli was significantly stronger in V1 compared to LGN (2-, 4- and 9-back stimulus, two-sided Welch’s unequal variances t-test, all p = 0.02) and LP (3-back stimulus, p = 0.001, corrected for multiple comparisons), even after accounting for differences in the initial strength of adaptation between thalamus and V1 (i.e. normalizing to the 1-back adaptation ratio). The temporal decay of adaptation in higher-level extrastriate areas was similar to the decay in V1 (**Figure 3B)** and long-term adaptation in cortical areas was very consistent across mice (**Figure 3C**, right).

We further characterized the timescale of recovery from adaptation in visual cortex and thalamus by fitting exponential decay models to the n-back adaptation ratios in the respective areas. Recovery in cortical visual areas was better explained by double-exponential decay models, with a fast and a slow-decaying adaptation component, compared to a single-exponential decay (F-tests, all p < 0.006, except VISrl: p = 0.072). Recovery from adaptation was slowest in V1 with an exponential time constant τ_slow_ of 6.82 trials (bootstrapped 95% CI [3.39, 13.50]; **Figure 3A**, black dashed line), but was relatively similar for extrastriate areas (τ_slow_ ranging from 3.12 to 5.52 trials, all 95% CI intervals overlapping, see **Table 1** for all parameter estimates). In contrast, the recovery of adaptation in the thalamus was most parsimoniously captured by a single-exponential decay model (F-tests, p = 1 for both LGN and LP), and the time constants of the single-exponential decays were very short (LGN: τ_fast_ = 0.02 trials, 95% CI [0.004, 0.63]; LP: τ_fast_ = 0.71 trials, 95% CI [0.03, 0.93]). Together, these results indicate that adaptation in response to relatively brief 2-second stimuli decays surprisingly slowly in cortical visual areas, over the course of dozens of seconds, and survives the presentation of multiple intervening stimuli. Conversely, while adapting to the immediate stimulus history, neurons in the thalamus exhibit a fast recovery from adaptation, in line with a shorter temporal integration timescale for low-compared to high-level visual areas.

**Table 1.** Best fitting parameters of exponential decay models fitted to adaptation ratios (drifting gratings). Amplitude parameters ***a*** are expressed in %-response reduction of *repeat* with respect to *orthogonal* trials. Exponential time constants τ are expressed in units of trials. The decay of adaptation in thalamic areas LGN and LP was significantly better fit by single-exponential decay models. Therefore, no parameters for the second exponential component are provided for these areas. Values in parentheses indicate bootstrapped 95% confidence intervals.

| ROI | $a_{\text{fast}}$ | $\tau_{\text{fast}}$ | $a_{\text{slow}}$ | $\tau_{\text{slow}}$ |
| --- | --- | --- | --- | --- |
| V1 | **13.99** [10.73, 16.14] | **0.85** [0.59, 1.13] | **3.45** [1.71, 6.64] | **6.82** [3.39, 13.50] |
| AL | **10.19** [8.14, 12.71] | **0.01** [1e-3, 0.47] | **6.84** [4.90, 8.54] | **4.08** [3.10, 5.94] |
| AM | **9.96** [6.88, 14.30] | **0.39** [5e-3, 0.87] | **8.03** [3.83, 11.04] | **3.39** [2.37, 6.50] |
| LM | **13.35** [10.37, 16.07] | **0.46** [0.02, 0.72] | **3.71** [1.89, 5.84] | **5.52** [3.42, 10.60] |
| PM | **13.20** [10.43, 16.32] | **0.02** [2e-3, 0.45] | **6.50** [3.91, 9.26] | **4.43** [2.75, 9.28] |
| RL | **4.58** [1.97, 12.00] | **0.26** [5e-3, 1.83] | **7.40** [0.7, 9.82] | **3.12** [2.19, 34.41] |
| LGN | **6.51** [4.04, 8.85] | **0.02** [4e-3, 0.63] | - | - |
| LP | **16.76** [12.68, 21.02] | **0.71** [0.03, 0.93] | - | - |

### Cortical long-term adaptation is due to suppression following stimulus repetitions

Thus far we have quantified adaptation as the ratio of neural responses following repeated versus orthogonal stimuli. While this quantification revealed orientation-specific long-term traces of past stimuli in visual cortex, they do not reveal the relative contribution of response suppression (when a past orientation is repeated), and response enhancement (when the current and past orientations are orthogonal). To assess this, we leveraged the presentation of randomly interspersed blank trials during which no stimulus was presented. In particular, we used trials for which the *past n-back stimulus was a blank trial* to establish a baseline adaptation effect against which to compare trials for which the n-back stimulus was repeated or orthogonal. We found that 1-back repeated and orthogonal stimulus presentations both suppressed neural response (**Fig. 5**). Importantly, the suppressive effect of orthogonal orientations decayed quickly, and was limited to the 1-back trial, whereas the suppressive effect of repeated stimuli decayed more slowly, and remained significant for up to 8 trials back (**Fig. 5**). This suggests that long-term adaptation effects are mainly driven by response suppression to repeated stimulus orientations.

### Long-term adaptation following exposure to brief static gratings

So far, we have shown that neurons in mouse visual cortex exhibit long-lived adaptation to 2-second presentations of drifting gratings, influencing subsequent visual processing over the time course of dozens of seconds and multiple intervening stimuli. However, it is unclear to which degree the existence of such long-term adaptation effects depends on the particular stimulus type (drifting gratings) and duration (2 seconds). We therefore tested whether similar long-lived adaptation effects can be elicited by the presentation of brief, static gratings. Mice were presented with a rapid stream of static gratings, presented back-to-back for 250 milliseconds each (**Figure 6A**). Similar to our previous analyses, we probed orientation-specific adaptation by contrasting visual responses to gratings that were preceded by a grating of the same or orthogonal orientation. In V1, the repetition of stimulus orientation led to a clear reduction of the visual response to the current grating (**Figure 6B**, green shaded area; *n* = 530; adaptation ratio: 0.90, p = 2e-81, 95% CI [0.89, 0.91]). Very similar adaptation effects were found for extrastriate areas (**Figure 6C**, 1-back adaptation ratios between 0.89 and 0.93, all p < 8e-9, two-sided t-tests, corrected for multiple comparisons) and adaptation was consistent across mice (**Figure 6D**). Although there were only relatively few responsive neurons in the thalamus (LGN: n = 60; LP: n = 16), both LGN and LP exhibited significant adaptation effects (LGN - adaptation ratio: 0.93, p = 0.004, 95% CI [0.93, 0.98]; LP – adaptation ratio: 0.85, p = 0.01, 95% CI [0.85, 0.96]). Overall, these findings of orientation-specific adaptation, exerted by the immediately preceding static grating stimulus, parallel those found for adaptation to drifting gratings.

Next, we investigated the timescale over which adaptation to briefly presented static gratings affected subsequent visual processing. Neurons in V1 showed significant adaptation effects to stimuli presented as far as 20 presentations (5 seconds) in the past (**Figure 7A** and **7C** showing consistent adaptation across mice). Again, the decay of adaptation was well described by a double exponential decay model with a long time constant τ_slow_ = 9.12 trials (95% CI [6.09, 14.82]; **Figure 7A**, black dashed line). Higher-level extrastriate cortical areas showed similar decay dynamics (**Figure 7B**), with decay time constants ranging from 6.54 (VISam) to 21.78 trials (VISpm; all 95% CI intervals overlapping; see **Table 2** for all parameter estimates). While all cortical areas were significantly better fit by a double exponential decay model (F-tests, all p < 1e-5), neurons in LGN were more parsimoniously described (p = 0.22) by a single exponential decay with a shorter time constant (τ_fast_ = 2.93 trials, 95% CI [1.36, 7.32]), replicating the experiment using drifting gratings. However, in this experiment, the difference in long-term adaptation of cortex and thalamus was less pronounced than in the experiment using drifting gratings. We did not include thalamic nucleus LP in this analysis due to the low number of visually response neurons in this area (16 neurons across 32 mice). These findings demonstrate that even briefly presented static grating stimuli, which are embedded in a rapid stream of stimulus presentations, still elicit robust long-term cortical adaptation effects that persist despite the encoding of many intervening stimuli.

**Figure 7.**
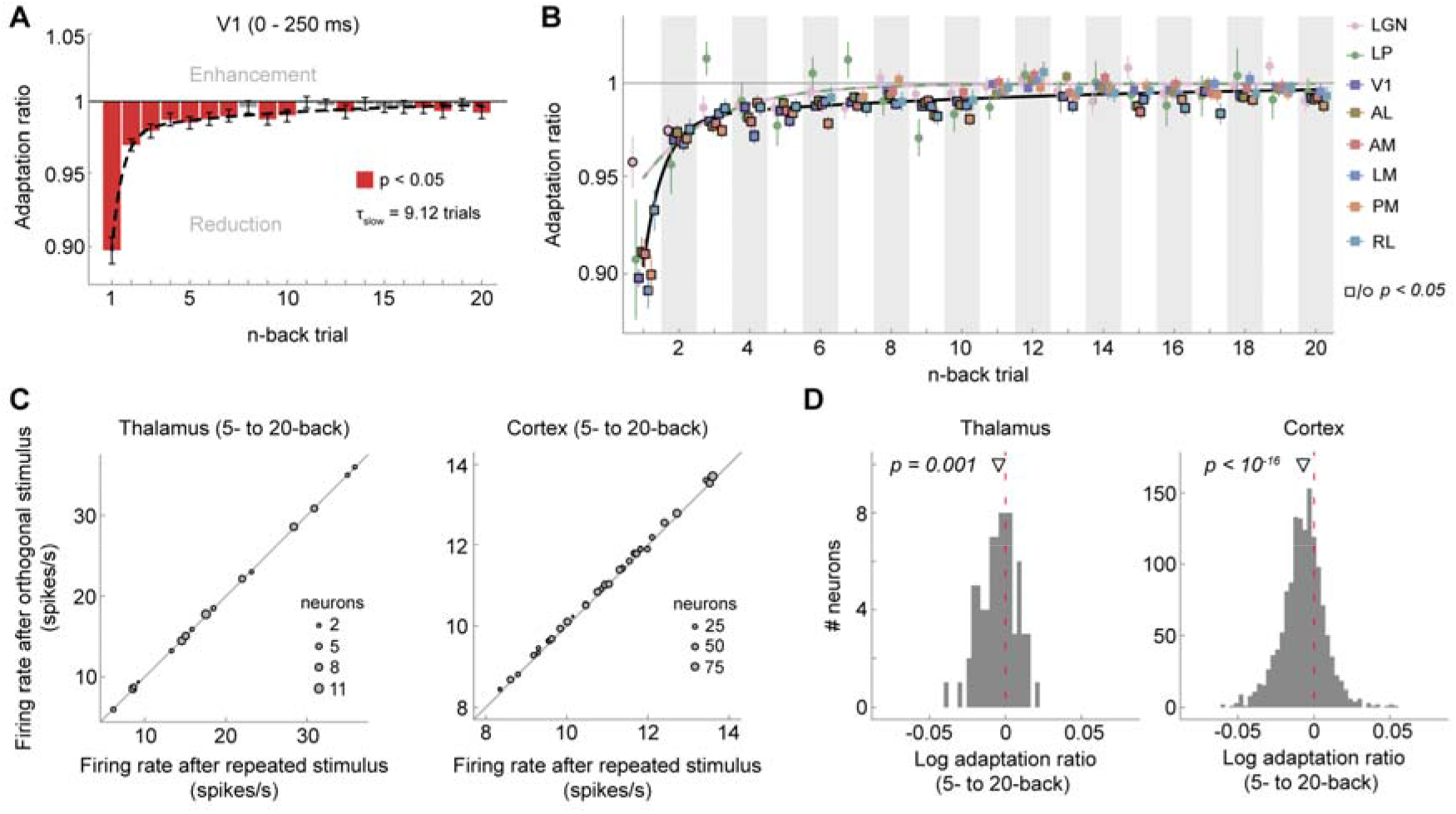
Visual cortex exhibits long-term adaptation following briefly presented gratings. **(A)** Adaptation ratios of V1 as a function of the n-back trial. While adaptation was most strongly driven by the previous stimulus (1-back), stimuli encountered up to 20 presentations in the past (5 seconds ago) still exerted significant adaptation effects on the current visual response (red bars, p < 0.05, FDR-corrected). Similar to drifting grating adaptation, the decay of adaptation over n-back trials was well captured by a double-exponential decay model with a fast- and slow-decaying adaptation component (black dashed line; *a_fast_* = 8.17%, t_fast_ = 0.54 trials, *a_slow_* = 2.04%, t_slow_ = 9.12 trials). Error bars denote bootstrapped 95% confidence intervals. **(B)** Adaptation ratios as function of n-back trial for different visual areas (color-coded). In cortical areas (square symbols) there is significant adaptation to stimulus orientations presented up to 20 trials back (symbols with black border, p < 0.05, FDR-corrected per area), while in thalamic areas (circle symbols) long-term adaptation is less evident. Error bars denote standard errors of the mean. Black and orange-green lines denote the best fitting exponential decay models for cortex and thalamus, respectively. Adaptation was computed over the whole stimulus interval (0 to 250 ms), since due to the back-to-back presentation of static gratings, visual responses to the previous stimulus overlapped with the initial time window of the current stimulus, thereby increasing response variability in this early time window. However, largely similar results were obtained when performing the analyses on the same time window used in the drifting grating experiment (0 to 100 ms), except for a less clear difference of the decay of adaptation between cortex and thalamus. **(C)** Average firing rates per mouse when the 5- to 20-back orientation was repeated (*x*-axis) or orthogonal (*y*-axis) relative to the current orientation, in the thalamus (left) and cortex (right). **(D)** Histograms of single-neuron long-term (avg. 5- to 20-back) adaptation ratios (log-transformed) in thalamus (left) and cortex (right).

**Table 2.** Best fitting parameters of exponential decay models fitted to adaptation ratios (static gratings). Amplitude parameters ***a*** are expressed in %-response reduction of *repeat* with respect to *orthogonal* trials. Exponential time constants τ are expressed in units of trials (250 ms duration). The decay of adaptation in LGN was significantly better fit by single-exponential decay model. Therefore, no parameters for the second exponential component are provided for LGN. Values in parentheses indicate bootstrapped 95% confidence intervals.

| ROI | $a_{fast}$ | $\tau_{fast}$ | $a_{slow}$ | $\tau_{slow}$ |
| --- | --- | --- | --- | --- |
| V1 | **8.17** [7.11, 9.23] | **0.54** [0.37, 0.71] | **2.04** [1.33, 2.91] | **9.12** [6.09, 14.82] |
| AL | **7.38** [5.27, 8.64] | **0.68** [0.02, 1.05] | **1.49** [0.77, 3.25] | **14.79** [5.24, 39.75] |
| AM | **6.24** [4.46, 8.05] | **0.48** [8e-3, 0.96] | **2.77** [1.13, 4.43] | **6.54** [3.82, 19.11] |
| LM | **7.93** [6.14, 9.80] | **0.40** [0.02, 0.61] | **2.91** [1.89, 4.00] | **7.74** [5.42, 13.36] |
| PM | **8.46** [6.74, 10.30] | **0.64** [0.36, 0.87] | **1.57** [1.10, 2.25] | **21.78** [13.07, 37.00] |
| RL | **5.39** [3.50, 7.56] | **0.66** [0.02, 1.09] | **1.39** [0.87, 2.07] | **19.58** [10.99, 37.51] |
| LGN | **3.91** [1.80, 6.59] | **2.93** [1.36, 7.32] | - | - |

### Short-term adaptation does not introduce spurious long-term adaptation effects

Our analysis approach of quantifying adaptation to the n-back stimulus by conditioning the current visual response on the orientation difference between current and previous n-back stimulus (repeat/orthogonal) relies on the assumption that the stimulus sequence is uncorrelated. If the presentations stimulus orientations were correlated across trials, these correlations may introduce spurious adaptation effects, potentially causing short-term adaptation to masquerade as long-term adaptation (Maus et al., 2013). While the presentation order of stimuli of the current experiments was randomized, making such spurious adaptation effects unlikely, we nevertheless assessed this potential confound via the simulation of an artificial neuron that only exhibited short-term (1-back) adaptation. We observed no spurious long-term adaptation effects for this artificial neuron when presented with the drifting grating sequences (**Figure 8A**), nor when presented with the static grating sequences (**Figure 8B**), markedly different from the long-term adaptation effects we observed in the empirical data.

**Figure 8.**
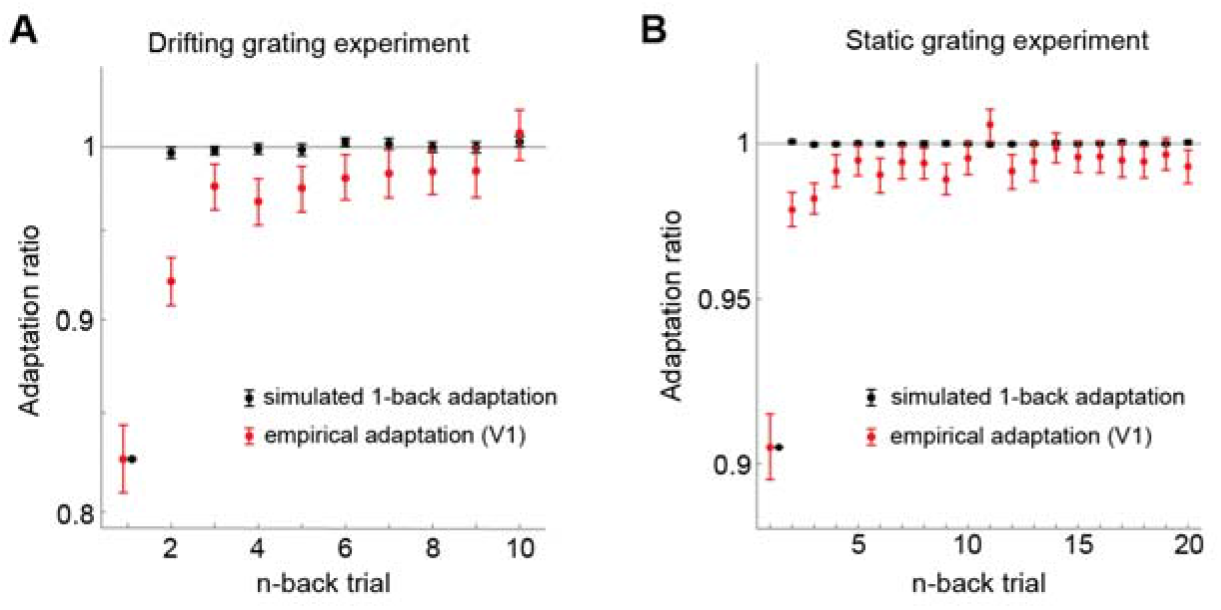
Short-term (1-back) adaptation does not introduce spurious long-term adaptation effects for the particular stimulus sequences used in the experiments. We simulated responses of a artificial neuron to the particular stimulus sequences used in the drifting grating experiment (**panel A**) and static grating experiment (**panel B**). The artificial neuron responded equally to all stimulus orientations, but selectively reduced its responses to a successive repeated orientation to mimic orientation-specific 1-back adaptation. We chose the strength of this 1-back adaptation effect to match the empirically observed 1-back adaptation of V1. We subsequently analyzed the simulated responses with the same procedure used for the empirical data. The analysis of the simulated responses recovered the ground truth 1-back adaptation effect (black data points). There were no spurious adaptation effects for stimuli further in the past, as indicated by the black data points being centered on an adaptation ratio of 1, markedly different from the empirically observed long-term adaptation effects (red data points - adaptation in V1). Black error bars denote 95% CIs of adaptation across the simulations of the 32 stimulus sequences. Red error bars denote 95% CIs of empirical adaptation across neurons in V1.

## Discussion

We observed that neurons in mouse visual cortex exhibit remarkably long timescales of adaptation effects after brief visual stimulation, influencing the processing of subsequent input over dozens of seconds and outliving the presentation of several intervening stimuli. The long-term adaptation effect was stimulus-specific - tuned to the orientation differences between past and current stimuli - indicating that the visual cortex maintains a lasting memory trace of individual briefly experienced stimuli. Although adaptation to individual past stimuli was subtle, the expected cumulative adaptation effect of the long-term stimulus history outweighed short-term adaptation to the immediately preceding stimulus. This suggests that long-term adaptation can have a profound influence on sensory processing, especially when visual input is temporally correlated, as is the case for natural environments (van Bergen and Jehee, 2019). While adaptation to drifting gratings decayed at a similar rate in primary and extrastriate visual cortex, and was still observable for stimuli seen 8 trials (or 22 seconds) in the past, adaptation in the thalamus decayed more quickly, limited to the 1- or 2-back stimulus (experienced 1-4 seconds prior). This demonstrates that the long-term component of adaptation observed in the visual cortex is not inherited from the thalamus, but is maintained in cortical circuits. Finally, we replicated our findings of cortical long-term adaptation to drifting gratings with a different stimulus set of rapidly presented static gratings, underlining the robustness and ecological validity of the long-term temporal dependencies. However, in this experiment, the difference in long-term adaptation of cortex and thalamus was less pronounced than in the experiment using drifting gratings. The back-to-back presentation of static gratings may interfere with our measurement of adaptation effects, because responses during stimulus presentation is likely to include both responses to the onset of that stimulus, and responses to the offset of the previous stimulus. Together, our findings show that visual cortex maintains concurrent stimulus-specific memory traces of briefly presented input, which allow the visual system to build up a statistical representation of the world over longer timescales. We speculate that this may enable the visual system to exploit temporal input regularities over extended timescales to efficiently encode new visual stimuli under natural conditions (Barlow and Földiák, 1989; Müller et al., 1999; Wainwright, 1999; Clifford et al., 2000; for reviews see Schwartz et al., 2007; Weber et al., 2019).

There is ample evidence that sensory cortex can exhibit long-term adaptation following long exposure to a stimulus. For instance, long stimulus presentations lasting from dozens of seconds to several minutes can alter visual responses of neurons in monkey and cat primary visual cortex over similarly long timescales, persisting for several minutes (Dragoi et al., 2000; Patterson et al., 2013). Furthermore, stimulus-specific adaptation effects can accumulate over many brief intermittent presentations of the same stimulus (Kuravi and Vogels, 2017), and subsequently show a persistence of several seconds (Ulanovsky, 2004; Peter et al., 2020). Crucially, in contrast to these previous studies, here we tested the adaptation effects elicited by individual, briefly presented stimuli. In the stimulus sequences of the current experiments, all stimulus orientations occurred equally often and in random order, precluding systematic accumulation of adaptation to any particular orientation of higher prevalence. Despite the absence of such accumulation effects, we find that the presentation of brief individual stimuli alters subsequent visual processing over time spans of at least 22 seconds and affects the processing of many subsequent stimuli. This demonstrates that long-term adaptation effects are not contingent on long adaptor durations or many repeated presentations of the same adaptor stimulus, but can occur in much more naturalistic settings that are also frequently employed in experimental designs, i.e. in response to brief individual visual experiences.

The observation of long-term adaptation effects to brief stimuli is particularly surprising, as previous studies investigating the recovery of adaptation following brief visual stimulation reported only very fleeting adaptation effects. In V1 of anesthetized monkeys, adaptation to 4-seconds long drifting gratings decayed with a half-life of ~1 second, in the absence of any intervening visual input (Patterson et al., 2013). This half-life is much shorter than the ~14 seconds we observed in the current study. We speculate that this difference could be, at least partly, related to the anesthetized versus awake state of the animals in the respective experiments, and that long-term adaptation might be facilitated by deeper, recurrent stimulus processing in awake animals. Nevertheless, recent studies in awake mice point towards similar short-lived adaptation effects in V1. For instance, adaptation to 100 ms gratings has resulted in decay time constants of 0.5 to 1 second (Jin et al., 2019; Jin and Glickfeld, 2020), and other studies have found no detectible effects of adaptation to 2-seconds drifting gratings after a 6-seconds delay (King and Crowder, 2018), and no history dependencies beyond 1 second in response to 250 milliseconds orientation patterns (Kim et al., 2019). Notably, most of these previous studies have investigated adaptation in the absence of any intervening visual input, making the current observation of stimulus specific long-term adaptation despite intervening input even more astounding. One major advantage of the current study is the large number of recorded neurons (2,365), which vastly increased our power to reveal subtle but reliable long-term adaptation effects that may have gone unnoticed in previous studies.

It should be noted that long-term adaptation, also in the face of intervening visual input, has been observed in higher-order visual areas in infero-temporal cortex of primates and humans, when observers performed a task on repeated stimuli (Henson et al., 2000, 2004; McMahon and Olson, 2007). Importantly, these long-term adaptation effects, also known as repetition suppression (Grill-Spector et al., 2006; Barron et al., 2016), appear to be highly dependent on attention (Murray and Wojciulik, 2004; Henson and Mouchlianitis, 2007; Larsson and Smith, 2012) and task (Henson et al., 2002; Henson, 2016) and have been related to processes of memory recall (Meyer and Rust, 2018). While phenomenologically similar to the current adaptation effects (reduction of neural activity), it is likely that these task-dependent higher-level repetition suppression effects are distinct from the automatic and early adaptation effects on sensory encoding measured in the current experiments, which take place in both primary and higher-order visual areas in the absence of an explicit task. In support of this view, previous studies measuring long-term repetition suppression effects in infero-temporal cortex in the presence of a task did not observe concomitant long-term effects in early visual cortex (Sayres and Grill-Spector, 2006; Weiner et al., 2010), suggesting that these high-level repetition suppression effects are at least partially distinct from automatic and early adaptation effects on sensory encoding. In contrast, here we show that even in the absence of an explicit task, the earliest stages of cortical visual processing automatically adapt to the long-term history of individual briefly presented stimuli.

It has been previously proposed that temporal integration timescales increase along the cortical hierarchy (Hasson et al., 2008; Lerner et al., 2011; Honey et al., 2012; Murray et al., 2014). Here, we show that the integration window of temporal context, in the form of adaptation, increases from the thalamus to cortex, broadly in line with these proposals. However, we did not find different integration times between lower-level primary and higher-level extrastriate visual cortex, congruent with a recent study in humans (Fritsche et al., 2020a; but see Zhou et al., 2018). Since we measured adaptation in the early feedforward response (0 to 100 milliseconds), it appears unlikely that long-term adaptation in V1 was inherited from higher-level visual areas through feedback connections, but rather suggests that long-term temporal context already influences the earliest stages of cortical processing. The similar decay of adaptation across cortical areas could either be due to the comparatively flat hierarchical structure of mouse visual cortex (for a review see Glickfeld and Olsen, 2017) or may reflect an important difference between the temporal tuning of adaptation and previously reported temporal integration timescales.

Importantly, while the current study focused on the early feedforward response (first 100 ms for drifting gratings, 250 ms for static gratings), adaptation has been found to alter neural responses beyond the early response epoch, further interacting with factors such as stimulus size and adaptation duration (Patterson et al., 2013), pointing towards more complex inhibitory and excitatory population-level coordination (Solomon and Kohn, 2014). In order to obtain a full understanding of the sources and the long-term consequences of adaptation, future studies will therefore need to investigate further properties of the long-term adaptation effects reported here, such as their dependence on stimulus parameters and response epoch.

Recent psychophysical studies in humans have revealed long-lived repulsive perceptual biases following briefly presented gratings, biasing subsequent orientation perception over dozens of seconds (Chopin and Mamassian, 2012; Suárez-Pinilla et al., 2018; Gekas et al., 2019; Fritsche et al., 2020b). Our current findings of long-term orientation-specific adaptation in early visual cortex suggests a potential neural mechanism underlying these perceptual biases. Interestingly, a recent behavioral study in rats revealed similar long-term dependencies in a vibrissal vibration judgment task (Hachen et al., 2020), suggesting potential parallels of long-term perceptual adaptation between rodents and humans. An important future goal will be to quantitatively relate such behavioral adaptation biases to the present long-term history dependencies at the neural level.

To conclude, our findings highlight the ubiquitous influence of the short- and long-term stimulus history on current sensory processing in visual cortex. This dependence on the broader temporal context may enable the visual system to efficiently represent information in a slowly changing environment (Schwartz et al., 2007; Weber et al., 2019).

